# NMR structure of the carboxy-terminal domain of the Ea22 pro-lysogenic protein from lambda bacteriophage

**DOI:** 10.1101/2023.09.01.555870

**Authors:** Cameron Goddard, Bożena Nejman-Faleńczyk, Logan W Donaldson

## Abstract

The *ea22* gene resides in a relatively uncharacterized region of the lambda bacteriophage genome between the *exo* and *xis* genes and is among the earliest genes transcribed upon infection. In lambda and Shiga toxin-producing phages found in enterohemorrhagic *E. coli* (EHEC) associated with food poisoning, Ea22 favors a lysogenic over lytic developmental state. The Ea22 protein may be considered in terms of three domains: a short amino-terminal domain, a coiled-coiled domain, and a carboxy-terminal domain (CTD). While the full-length protein is tetrameric, the CTD is dimeric when expressed individually. Here, we report the NMR solution structure of the Ea22 CTD that is described by a mixed alpha-beta fold with a dimer interface reinforced by salt bridges. A conserved mobile loop may serve as a ligand for an unknown host protein that works with Ea22 to promote bacterial survival and the formation of new lysogens. From sequence and structural comparisons, the CTD distinguishes lambda Ea22 from homologs encoded by Shiga toxin-producing bacteriophages.

## Introduction

Viral dark matter is a term given to describe the vast number of unknown genes and their products^1^. Even bacteriophage λ, whose history spans the entirety of molecular biology, has a genetic region between the *exo* and *xis* genes that has remained under-characterized, until recently. Driven by the *p*_L_ promoter, the λ *exo-xis* region genes comprising *orf61, orf63, orf60a, orf73*, and *ea22* are among the earliest transcribed during infection^2^ (Fig. 1). The *exo-xis* region was associated with cell cycle synchronization^3^ and inhibition of host DNA replication^4^. Since these initial observations, *exo-xis* region genes have been identified as regulators of the viral life cycle and the decision to either actively produce progeny and lyse the host cell (lytic cycle) or remain domain and integrated into the host genome (lysogenic cycle). Since deletion of the *exo-xis* region does not inhibit replication^4^, it leads to the idea that *exo-xis* region genes serve an auxiliary and more nuanced role during development possibly by engaging host transcription factors, toxin-antitoxin regulatory pathways, and stress response pathways.

**Figure 1.**
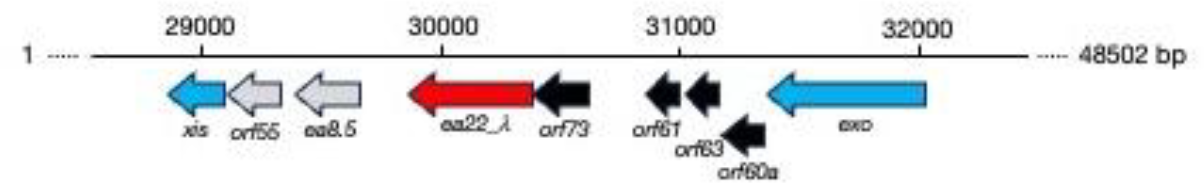
The *exo-xis* region of phage λ. A genomic cassette including ea22 (red) and four other genes (black) are typically observed among lambdoid phages. Beyond this cassette, these is considerable diversity. In phage λ, two additional genes are observed (grey).

At 182 amino acids in length, Ea22 is the largest of the *exo-xis* proteins. A combination of sequence comparisons, structure predictions, and deletion studies suggest that Ea22 may be considered in three parts consisting of a short variable N-terminal region, a central coiled-coiled region comprising over half the protein, and a C-terminal region. While full-length λ Ea22 is tetrameric in solution; the C-terminal domain (CTD) alone is dimeric suggesting that the functional organization of λ Ea22 may be considered as a dimer-of-dimers^5^. Building upon these molecular biological and biochemical studies, the NMR structure of the λ Ea22 CTD is presented in this report. The Ea22 CTD structure represents the second high-resolution examination of the *exo-xis* region, the first being the NMR structures of two λ Ea8.5 homologs^6^. A recent review of all *exo-xis* region proteins predicted by AlphaFold and RoseTTAfold machine learning methods includes a general discussion of Ea22^7^.

While the shorter and presumably single domain *exo-xis* proteins Orf61, Orf63 and Orf60a promote the lytic developmental pathway^8,9^, Ea22 remains in sole contrast as pro-lysogenic developmental protein^10^. Providing evidence for this role, bacterial lysogens with a deleted *ea22* gene are rapidly induced towards lytic development by UV irradiation and produce more viral progeny as the infection proceeds^10^. Since all *exo-xis* region proteins are expressed within minutes of infection, their combined action towards promoting lytic or lysogenic development may be the net result of several host metabolic and stress related pathways being interrogated or manipulated at once.

Enterohemorrhagic *E. coli* (EHEC) such as O157:H7^11–13^ are responsible for food poisoning outbreaks and more severe outcomes in immunocompromised people^14^. These strains contain a variety of genetic elements that contribute to their pathogenicity, including being lysogens of phages that share many similarities with phage λ^15^. One important distinction between the resident prophage sequences within these strains and phage λ is the presence of a gene encoding Shiga toxin (either *stx1 or stx2*) that is produced during the late stages of the lytic developmental cycle^16^. Like ricin, Shiga toxin disables ribosomes in intestinal epithelial cells thus aggravating the bacterial infection. Stx phages also contain copies of *exo-xis* genes. Differences in gene expression and developmental effects arising from *exo-xis* region gene expression in Stx phages tend to be more amplified^17,18^. The *ea22* gene is most illustrative of this distinction between Stx phages and phage λ since not only are the effects more pronounced, but the Ea22 proteins of Stx phages have CTDs that are distinct from the Ea22 CTD of phage λ.

The λ Ea22 CTD alone cannot reproduce the pro-lysogenic properties of the full-length protein^5^ suggesting that the CTD does not have any intrinsic activity or it requires support from the amino-terminal sequence and central coiled-coil domain. From previously published multi-angle laser scattering studies, the λ Ea22 CTD was observed to be exclusively dimeric^5^. The full-length protein; however, was observed to be exclusively tetrameric^5^. It is currently unknown if the coiled-coil domain mediates tetramerization or the protein is organized as dimer-of-dimers via interactions contributed by the short terminal amino-terminal segment. A previous comparison of ^1^H,^15^N-HSQC spectra of full-length λ Ea22 and the λ Ea22 CTD revealed similar spectra^5^ suggesting the CTD was decoupled from the rest of the 84 kDa tetramer by a mobile segment, otherwise its ^1^H-^15^N resonances would have experienced the same degree of line broadening. During the same study, circular dichroism and differential scanning calorimetry revealed the Ea22 CTD was shown be soluble and thermostable in excess of 95 °C suggesting it would be an excellent candidate for high-resolution structural studies.

## Results

The NMR structure of the λ Ea22 CTD dimer was solved using a conventional heteronuclear approach that incorporated 993 intramolecular distance restraints 25 intermolecular distance restraints, 23 inferred hydrogen bonds, and 43 backbone dihedral angle restraint pairs that were predicted from chemical shifts. A complete statistical summary of the ensemble is available in Supplementary Table S1. The ensemble of the twenty best structure solutions (Fig. 2a) has a precision of 0.6 Å for ordered backbone atoms and 1.0 Å for all ordered heavy atoms.

**Figure 2.**
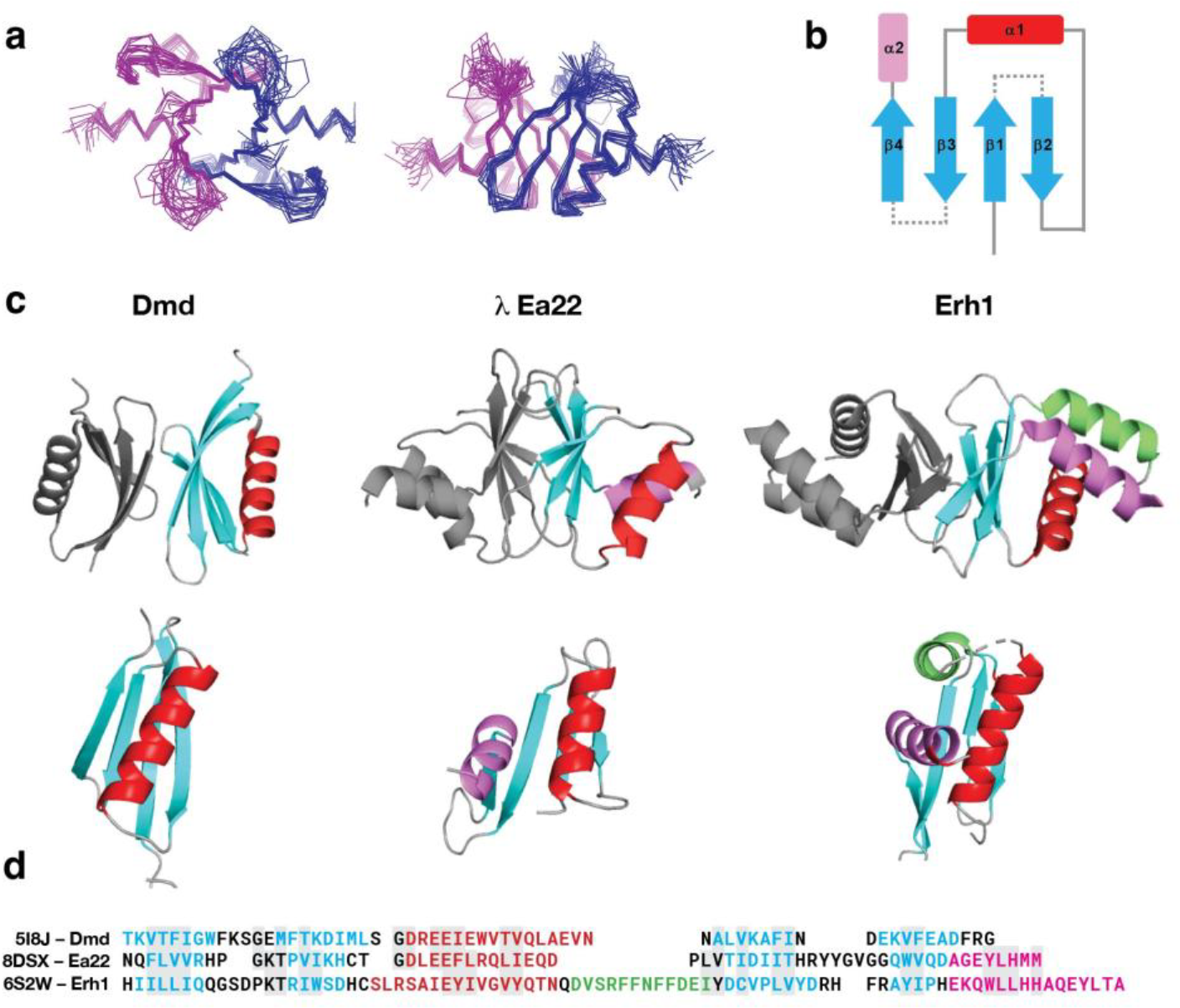
The NMR structure of the λ Ea22 CTD. (a) Two views of the structure ensemble with the two chains colored blue and magenta. **(b)** A schematic representation. **(c)** Similar dimeric proteins identified from the Protein Data Bank. To guide the comparison, topologically similar helices are colored red, purple and green. **(d)** A sequence alignment following the same coloring scheme of secondary structure elements.

The λ Ea22 CTD has a secondary structure composition of β1β2α1β3β4α1 (Fig. 2b). The two α-helices cross each other on one face of the four stranded β-sheet leaving the opposite face of the β-sheet to form the dimerization interface. The dimer interface incorporates contributions from all four β-strands (β1: F112/V114/R116; β2: P122/ I124; β3: T146/D148/I150; β4: V162). An ionic bond between R116 and D148 bridges the two protomers. An assigned ^1^H-^15^N HSQC spectrum is provided as Supplementary Fig. S1. Experimental data supporting the configuration of the β-sheet and a set of inter- and intramolecular NOEs supporting key contacts made by I124 is provided as Supplementary Fig. S2. A detailed view of the dimer interface is provided as Supplementary Fig. S3. Complete chemical shift assignments could not be made for the amino-terminal affinity tag and residues S102-Q111, the β2/α1 loop (H126-D130), and a longer β3/β4 loop (T151-G158) indicating a high degree of conformational freedom and motions in the µs-ms time scale.

The λ Ea22 CTD represents an example of how caution should be taken with machine learning predictions. Using the CTD sequence alone, the AlphaFold^19^ prediction of the dimer was very similar to the NMR structure (1.25 Å RMSD; Supplementary Fig. S4), with the only difference being that the AlphaFold method suggested a longer α2 helix that extending implausibly beyond the domain. In contrast, when the full-length sequence was used or a portion of the coiled-coil region was appended to the CTD, a completely different model was predicted. This model placed the α1 and α2 on opposing sides of the β-sheet rather than interacting together on the same side of the β-sheet. ESMfold is a complementary machine-learning based method to AlphaFold that uses a protein language model in lieu of a multiple sequence alignment^20^. The NMR solution structure was dissimilar to the ESMFold predicted models of the full-length sequences and the CTD sequence alone.

A structure-based search of the Ea22 CTD dimer against the PDB using FoldSeek^21^ and SSM^22^ did not identify any homologous structures. To present this unique configuration in context, the Ea22 CTD with its two α-helices is presented in Fig. 2c in between dimeric proteins that have either one or three α-helices. A sequence comparison follows the structural comparison in Fig 2d. The λ Ea22 CTD superimposes somewhat (2.39 Å RMSD) to the antitoxin protein Dmd from phage T4. From molecular docking experiments (not presented), the Ea22 CTD makes a reasonable fit with either RnlA or LsoA, two host toxin proteins that are inhibited by T4 Dmd to stop an infection-induced stress response. Taking the comparison further, the shared helix between the Ea22 CTD and T4 Dmd is positioned to make two key ionic and hydrogen bond contacts (Dmd:LsoA D28:R243 and Q35:R305 being analogous to Ea22:LsoA E38:R243 and E46:R305, respectively); however, a critical hydrophobic contact made by W34 and a secondary contact made by E9 in T4 Dmd are lost in the λ Ea22 CTD (E133 and E140, respectively). Given these important differences, Ea22 CTD would not fulfill the role of an antitoxin to *E. coli* LsoA and RnlA. The λ Ea22 CTD dimer has some superficial resemblance to *C. elegans* Erh1 which participates in a complex with three other proteins to facilitate the processing of regulatory RNAs^23^; however, not only is the orientation of the β-sheets is reversed and not as extensive, but there is also no analogous three-helix platform for supporting a donated fourth helix from a protein partner in the complex. In summary, given the lack of close homologs, the Ea22 CTD represents a unique structural assembly whose function remains unknown.

To investigate where variation is occurring in the Ea22 CTD, a search was performed against the four million non-redundant sequences comprising the UniRef100 database. From the fifty-four sequences identified spanning the entirety of the CTD (Supplementary Fig S5) amino acids with at least 30% variation were mapped on the λ Ea22 CTD NMR structure. As shown in Fig. 3, the most variation was observed at the solvent-facing sides of helices α1 and α2 and the β2/α1 loop. No variation was observed for the unstructured β3/β4 loop.

**Figure 3.**
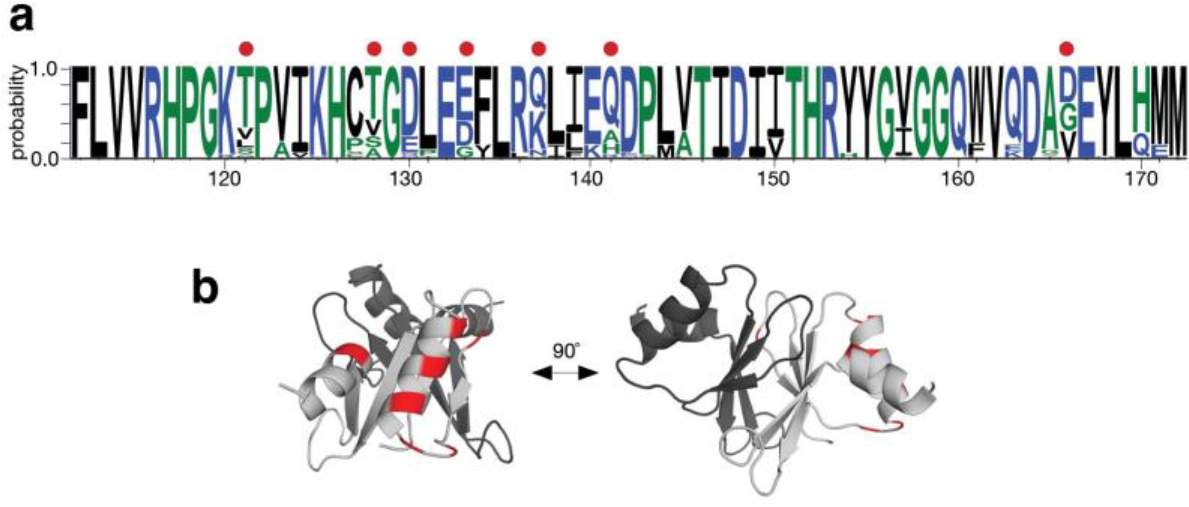
Sequence conservation in the λ Ea22 CTD protein fragments from the UniRef100 database. **(a)** WebLogo plot. A red dot indicates a position where >30% variability is observed. **(b)** The variability is mapped onto one chain (light grey) of the λ Ea22 CTD dimer.

The *ea22* gene is the last and largest of four conserved genes in the *exo-xis* region. In a standard laboratory *E. coli* MG1655 strain infected with any of four commonly studied Stx phages, *ea22* transcription exceeds all other *exo-xis* genes and early infection markers such as N^10^. However, in the same host strain infected with λ, the opposite result is observed with transcription being the lowest among the *exo-xis* region genes^10^. In addition to this major difference in gene expression, Stx phage *ea22* genes are mosaic^7,10^. To gain a greater understanding of variation among ea22 genes, a previously published dataset of thirty-seven phage and EHEC prophage genomes^24^ were analyzed. In eleven prophage genomes, no *ea22* protein was detected using a translated nucleotide BLAST search. The remaining genomes resembled λ *ea22* throughout the first half of the gene but no EHEC-associated *ea22* gene encoded a domain like the λ Ea22 CTD. Among the EHEC-associated *ea22* genes in the dataset, the observed Ea22 protein sequences could be reduced to three possible variants (Table 1). One sequence of each variant was selected as the prototype (Fig. 4). In eight cases. two Ea22 variants were encoded in the same genome. All three Ea22 variants were encoded in the genome of *E. coli* O157:H7 Xuzhou21, a strain that was first isolated in 1986 from China and was responsible for a large outbreak in 1999^25^ (Supplementary Fig. S6).

**Table 1.**
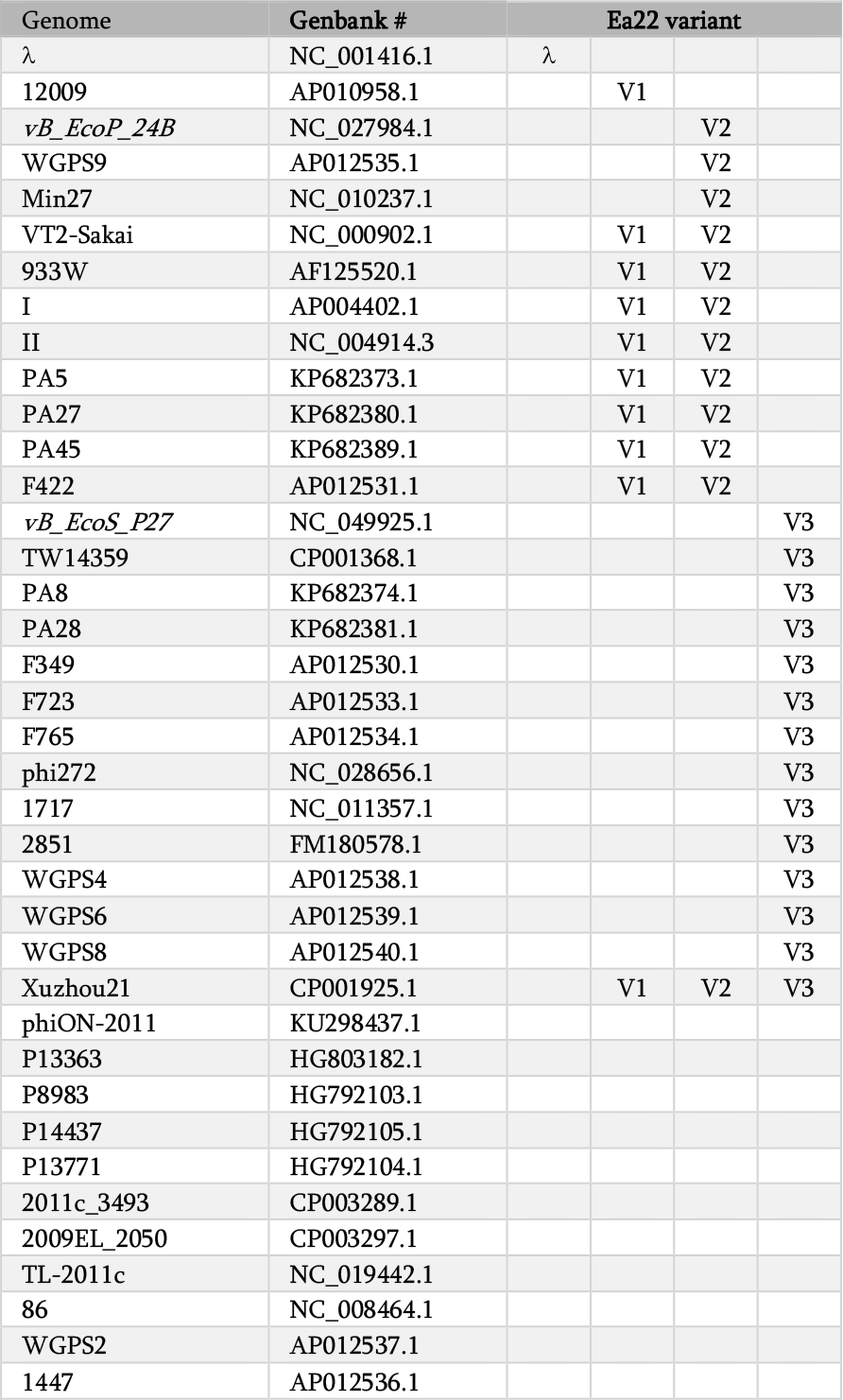
Classification of Ea22 variants in a collection of shigatoxigenic phages (italics) and EHEC strains.

| Genome | Genbank # | Ea22 variant |  |  |  |
| --- | --- | --- | --- | --- | --- |
| *λ* | NC_001416.1 | *λ* |  |  |  |
| 12009 | AP010958.1 |  | V1 |  |  |
| *vB_EcoP_24B* | NC_027984.1 |  |  | V2 |  |
| WGSP9 | AP012535.1 |  |  | V2 |  |
| Min27 | NC_010237.1 |  |  | V2 |  |
| VT2-Sakai | NC_000902.1 |  | V1 | V2 |  |
| 933W | AF125520.1 |  | V1 | V2 |  |
| I | AP004402.1 |  | V1 | V2 |  |
| II | NC_004914.3 |  | V1 | V2 |  |
| PA5 | KP682373.1 |  | V1 | V2 |  |
| PA27 | KP682380.1 |  | V1 | V2 |  |
| PA45 | KP682389.1 |  | V1 | V2 |  |
| F422 | AP012531.1 |  | V1 | V2 |  |
| *vB_EcoS_P27* | NC_049925.1 |  |  |  | V3 |
| TW14359 | CP001368.1 |  |  |  | V3 |
| PA8 | KP682374.1 |  |  |  | V3 |
| PA28 | KP682381.1 |  |  |  | V3 |
| F349 | AP012530.1 |  |  |  | V3 |
| F723 | AP012533.1 |  |  |  | V3 |
| F765 | AP012534.1 |  |  |  | V3 |
| phi272 | NC_028656.1 |  |  |  | V3 |
| 1717 | NC_011357.1 |  |  |  | V3 |
| 2851 | FM180578.1 |  |  |  | V3 |
| WGSP4 | AP012538.1 |  |  |  | V3 |
| WGSP6 | AP012539.1 |  |  |  | V3 |
| WGSP8 | AP012540.1 |  |  |  | V3 |
| Xuzhou21 | CP001925.1 |  | V1 | V2 | V3 |
| phiON-2011 | KU298437.1 |  |  |  |  |
| P13363 | HG803182.1 |  |  |  |  |
| P8983 | HG792103.1 |  |  |  |  |
| P14437 | HG792105.1 |  |  |  |  |
| P13771 | HG792104.1 |  |  |  |  |
| 2011c_3493 | CP003289.1 |  |  |  |  |
| 2009EL_2050 | CP003297.1 |  |  |  |  |
| TL-2011c | NC_019442.1 |  |  |  |  |
| 86 | NC_008464.1 |  |  |  |  |
| WGSP2 | AP012537.1 |  |  |  |  |
| 1447 | AP012536.1 |  |  |  |  |

**Figure 4.**
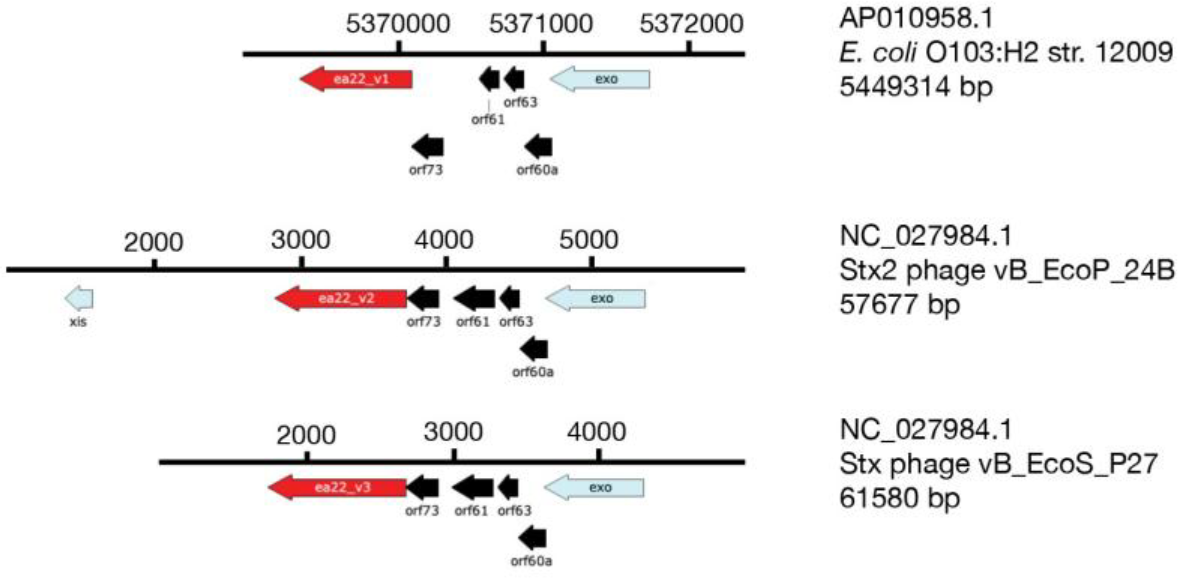
Three major variants of Ea22 are observed in a set of *Stx* phage and EHEC genomes. The *exo-xis* regions of an EHEC representative bearing *ea22* gene variant 1 (*ea22*_*v1*) and two Stx phage representatives bearing *ea22*_*v2* and *ea22*_*v3*, are shown. The encoded Ea22_V1 and Ea22_V2 and Ea22_V3 proteins serve as prototypes for further analyses.

A complete sequence comparison of λ Ea22 and the three Ea22 variants is presented as Supplementary Fig. S7. When the common regions are presented as blocks, the mosaic nature of the genes becomes apparent (Fig. 5a). Each variant shares the same amino-terminal segment and central coiled-coil region (with InterPro signature IPR025153/Ead_Ea22). The three *Stx* Ea22 variants contain a 63 amino segment that is absent in the λ Ea22 sequence. The carboxy-terminal sequences bear no similarity to each other except for a 16 amino acid segment in V1 and V2. An InterPro signature (IPR007539/DUF551) is unique to the Ea22 V2 variant. Like the λ Ea22 CTD, the DUF551 signature sequence has the hallmarks of a portable domain since it is found in *E. coli* RNA polymerase σ^70^, helix-turn-helix transcription factors, a cryptic prophage CPS-53 protein YfdS that alters sensitivity to oxidative stress^26^, and a dATP/dGTP pyrophosphohydrase from phage MazZ that inhibits host restriction enzymes^27^.

**Figure 5.**
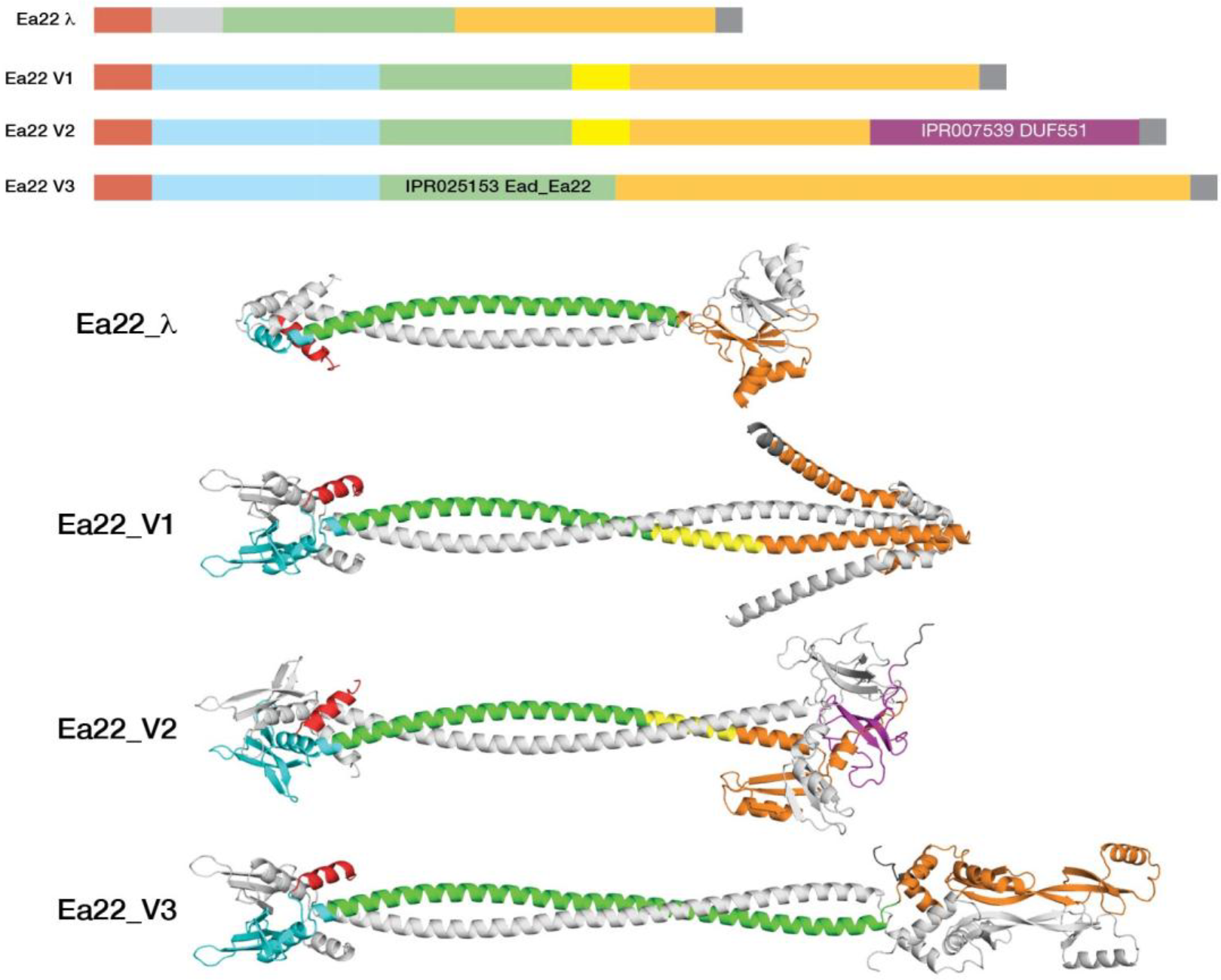
Ea22 proteins have a mosaic organization. All models were predicted by AlphaFold except for the λ Ea22 CTD which was grafted onto the full-length λ Ea22 protein. The green- and magenta-colored regions are signature sequences in the InterPro database. The N-terminal regions may be considered in terms of a common region shared with λ Ea22 (red) and another region (blue) only observed in the Ea22 variants. The CTD regions of each Ea22 (orange) are dissimilar.

While biochemical studies have determined that full-length λ Ea22 and Ea22 V3 (from phage *vB_ecoS_P27*) are tetrameric in solution, it is not presently known how the tetramer is formed. The two most straightforward models of a tetramer would be a dimer-of-dimers mediated by either the N-terminal region or the coiled-coil domain. Attempts to model the N-terminal and coiled-coiled regions as tetramers with AlphaFold Multimer produced candidates with poor packing. Given these modeling challenges, we took a reductionist rationale and modeled each Ea22 variant as a dimer using AlphaFold multimer (Fig. 5b). In Supplementary Fig. S8, the same modeling is presented using ESMFold that made similar predictions for the N-terminal and coiled regions. For the Ea22 V3 variant, AlphaFold and ESMFold diverged considerably in the predictions of the C-terminal regions. Neither prediction of the C-terminal region could explain why a stable fragment of the Ea22 V3 variant (termed P27 Ea22 in that study) was observed from limited proteolysis assay^5^.

## Discussion

The structure of the Ea22 CTD is unique to lambdoid phages. While the structure of the Ea22 CTD did not reveal clues towards its function in λ phage, its conservation suggests the preservation of a useful function in viral development. Shigatoxigenic phages and prophages associated with EHEC possess an *ea22* gene within the same *exo-xis* region of their genomes, albeit with some important structural differences. The most variability among the Ea22 proteins occurs at the C-terminal region of the protein. In a few cases where purified proteins can be studied, the C-terminal region is dimeric like λ regardless of sequence, and the full-length protein is tetrameric suggesting there is evolutionary pressure to maintain a specific oligomeric state even when the sequence and structure are drifting^5^. It is possible, therefore, that *Stx* phages have acquired new domains that improve fitness while maintaining a similar architecture. Within the complex environment of the human gut where bacteria experience oxidative attack by the immune system, Ea22 attenuates the lytic cycle thereby improving the number of bacterial survivors and those that become lysogens^10^. Interestingly, the uncharacterized motif (DUF551) of the Ea22_V2 is also observed in the protein YfdS that enhances resistance to oxidative stress^26,28^. Cells lacking *yfdS* lose the ability to adapt to their environment and are sensitive to H_2_O_2_ which is a commonly known natural inductor of Stx prophages in the human gut^18^. Importantly, the genes encoding Shiga toxins remain inactive within lysogenic bacteria, and the activation of prophages is required for their effective expression and the subsequent release of toxins. While UV light exposure or antibiotics that disrupt DNA replication are frequently employed in lab settings to trigger lambdoid prophages, such circumstances are unlikely to occur in the human intestinal environment. Exposure to H_2_O_2_ by intestinal neutrophils or protist predators creates oxidative stress that promotes the activation of Shiga toxin-converting prophages^17^. Thus, a potential role of Ea22 as a silencer of oxidative stress mediated Stx prophage induction is reasonable and may explain its pro-lysogenic activity^10^. The metabolic and signaling pathways through which lysogeny is maintained are likely employed by other stress responses since H_2_O_2_ induction as an effect is lower than UV irradiation or application of the DNA crosslinking antibiotic, mitomycin C.

Phage-phage and phage-host yeast two-hybrid surveys did not identify any potential partners of λ Ea22; however, these findings might have been influenced by factors such as the limited expression of one of the partners, interactions of low strength, and a requirement of Ea22 to function as one part of a multi-protein assembly^29,30^. If Ea22 accesses other phage and host proteins via the CTD, the β3/β4 loop (152-HRYYGVGG-159) serves as an interesting possibility since it is dynamic and can make both ionic and hydrophobic contacts.

Viral non-coding small RNAs (sRNAs) are established mediates of the development switch from the lysogenic to lytic development^31,32^. In the prevailing model, high concentrations of sRNAs act as a repressor of lytic gene expression. Since Ea22 is the only protein expressed from the *exo-xis* gene region that has pro-lysogenic characteristics^10^, we can only speculate at this time if it can act as a sRNA binding protein as a facilitator or mediator.

New structure prediction and sequence comparative methods offer a powerful means to jumpstart molecular and structural investigations of the vast amount of viral dark matter remaining to be explored. Future studies of proteins encoded by the *exo-xis* gene region may also reveal new ways in which to control the lysogenic-lytic developmental decision and make therapeutics more effective against food poisoning and a fraction of cases that become potentially life-threatening.

## Methods

### Expression and purification

A carboxy-terminal domain (CTD) fragment of λ Ea22 (102-182, reference #179655, UniProt: P03756) was gene synthesized by ATUM (Menlo Park, CA) and inserted into plasmid pD441NH for expression at 25 °C in an *E. coli* BL21:DE3 strain. At an A6_00nm_ of 0.8, expression was induced by the addition of 1 mM isopropyl thiogalactoside (IPTG) and the culture was grown for three hours further before harvesting. The expressed protein additionally contains a 6xHis tag at the amino-terminus and the sequence IETAV at the carboxy-terminus. Detailed information regarding the expression and purification of this fragment by nickel affinity and gel filtration chromatography has been published previously^5^. For NMR spectroscopy, a uniformly ^13^C, ^15^N labelled protein was made from a 1 L culture containing M9 minimal media salts supplemented with 3 g of ^13^C-glucose (Sigma-Aldrich), 1 g of ^15^N ammonium chloride (Sigma-Aldrich) and 1 g of ^13^C,^15^N Celtone algal extract (Cambridge Isotopes). Protein concentration was estimated by UV absorbance at 280 nm.

### NMR sample preparation and NMR spectroscopy

The λ Ea22 CTD purified protein was concentrated to 0.6 mM in 5 mM Tris-Cl, 50 mM NaCl, 0.05% (*w/v*) NaN_3_ and 10% (*v/v*) D_2_O for NMR spectroscopy. For NMR experiments requiring a sample in D_2_O, the Ea22 CTD protein was dialyzed to a similar buffer in 98% (*v/v*) D_2_O. A mixed ^12^C/^13^C sample was made by mixing ^12^C/^14^N and ^13^C/^15^N proteins at a 1:1 molar ratio, adding urea to 6 M and rapidly diluting the mixture into a 20-fold excess of NMR buffer. The protein was concentrated and dialyzed to a 98% (*v/v*) D_2_O buffer. A standard set of 2D and 3D heteronuclear NMR spectra were acquired at a temperature of 308 K on a 700 MHz Bruker AvanceIII spectrometer equipped with a nitrogen chilled probe. For backbone and side chain assignments, these experiments included (2D-15N-HSQC, 2D-13C-HSQC, 3D-HNCACB, 3D-CBCACONH, 3D-HNCA, 3D-HNCOCA, 3D-HNCO, 3D-HNCACO, 3D-CCONH, 3D-HCCONH, 3D-HBHACONH, and 3D-CCH-TOCSY). The 3D experiments were acquired according to a 10-20% sparse sampling schedule and processed with NMRPipe^33^ and HMSist^34^. Distance restraints were obtained from a 3D-15N-NOESY and a 3D-13C-NOESY experiments sparsely sampled at 25%. Intermolecular distance restraints were obtained from a 3D-^12^C-filtered/^13^C-edited NOESY experiment acquired at the laboratory of Lewis Kay (Univ. Toronto) on a Varian Inova 600 MHz instrument equipped with a room temperature probe.

### Structure determination

NMR spectra were analyzed with CCPN Analysis 2.52^35^. Backbone dihedral angles were predicted from chemical shifts with TALOS^36^. An initial ensemble of 100 structures sorted by the lowest number of NOE violations were calculated from a set of 10000 structures using CYANA 3^37^. The ensemble was refined using Rosetta build 2021.16.61629 with distance, angle and hydrogen bond restraints converted to Rosetta .cst format. Symmetry was enforced throughout the calculation. Details of the Rosetta refinement method have been published^38^. The top 20 refined structure solutions that satisfied the experimental restraints the most formed the final ensemble. Structural quality was assessed with PSVS^39^ and PROCHECK^40^.

### Bioinformatics

The structures of dimeric T4 Dmd (PDB: 5I8J) and the λ Ea22 CTD were superimposed with Superpose^22^ to identify analogous amino acids on Ea22 that would interact *E. coli* RnlA (PDB:6Y2P), LsoA (PDB:5HY3). The UniRef100 database^41^ was searched with mmseqs2^42^ for homologs to the λ Ea22 CTD. A few of the identified sequences in the initial set that were truncated beyond the minimally folded domain were removed. The dataset was realigned in AliView^43^ and exported as FASTA formatted list to WebLogo3^44^. The *exo-xis* regions of two phage genomes and a set of previously published prophage genomes and were mapped with TBLASTN using λ Exo, Xis, Orf60a, Orf61, Orf63, Orf55, and Ea22 proteins as query sequences. Identified genes were manually inspected and presented SnapGene (www.snapgene.com). AlphaFold Multimer was used to predict the dimeric structure of full-length λ Ea22 and one representative of each variant class described (https://github.com/sokrypton/ColabFold). A final full-length λ Ea22 dimer was constructed by using SWISS-MODEL^45^ to graft the N-terminal and coiled-coiled domains from the AlphaFold Multimer prediction to the NMR structure of the λ Ea22 CTD. Coiled-coiled region predictions were also made with CoCoPred^46^.

## Acknowledgments

This work was supported by a Canadian NSERC Discovery Grant (2018-05838) to L.W.D. and by National Science Center (Poland) within a project grant (UMO-2018/30/E/NZ1/00400) to B.N.-F.

## Author contributions statement

LWD conceived the experiments, performed the sequence analyses, acquired the NMR data, and solved the NMR structure. C.G. made methyl assignments of Ea22 CTD, performed molecular modeling, and helped manufacture protein samples. L.W.D wrote the manuscript, B.N.-F. and C.G. reviewed the manuscript.

## Additional information

### Data deposition

Chemical shifts of the λ Ea22 CTD were deposited in the BMRB (entry 51520). The final ensemble of structures was deposited in the PDB (entry 8DSX).

### Supplementary information

A separate datafile includes Table S1 and Figures S1-S8.

