## Supplementary material for "NMR structure of the carboxy-terminal domain of the Ea22 pro-lysogenic protein from lambda bacteriophage": Ea22-supplementary-information.pdf

**Table S1.** Statistics for the λ Ea22 CTD ensemble of structures

| | |
|---|---|
| NOE distance restraints in the ensemble[a] | 1018 |
| intraresidue | 493 |
| short (\|*i*-*j*\| = 1) | 182 |
| medium (1 ≤ \|*i*-*j*\| ≤ 5) | 93 |
| long (\|*i*-*j*\| > 5) | 225 |
| interchain | 25 |
| Hydrogen bond distance restraints | |
| HN–O / N–O pairs | 23 |
| Torsion angle restraints | |
| backbone (Φ / Ψ) | 43 |
| Structural quality analysis | |
| close contacts | 25 |
| RMS deviation of bond angles (deg) | 0.4 |
| RMS deviation of bond lengths (Å) | 0.009 |
| RMS deviation to the mean coordinates[b] | |
| all backbone / heavy atoms (Å) | 1.3 / 1.9 |
| ordered backbone / heavy atoms (Å) | 0.6 / 1.0 |
| Ramachandran plot[c] (%) | |
| residues in most favored regions | 96.1 |
| residues in additional allowed regions | 3.8 |
| residues in generously allowed regions | 0.1 |
| residues in disallowed regions | 0.0 |

[a] None of the twenty structures in the ensemble (PDB: 8DSX) has a distance violation > 0.2 Å and a dihedral angle violation > 5°.

[b] Ordered residues (113-118, 121-126, 130-151, 160-171) are defined by a dihedral angle order parameter with S(Φ)+S(Ψ)>=1.8 as determined by PSVS.

[c] Determined by PROCHECK for all residues.

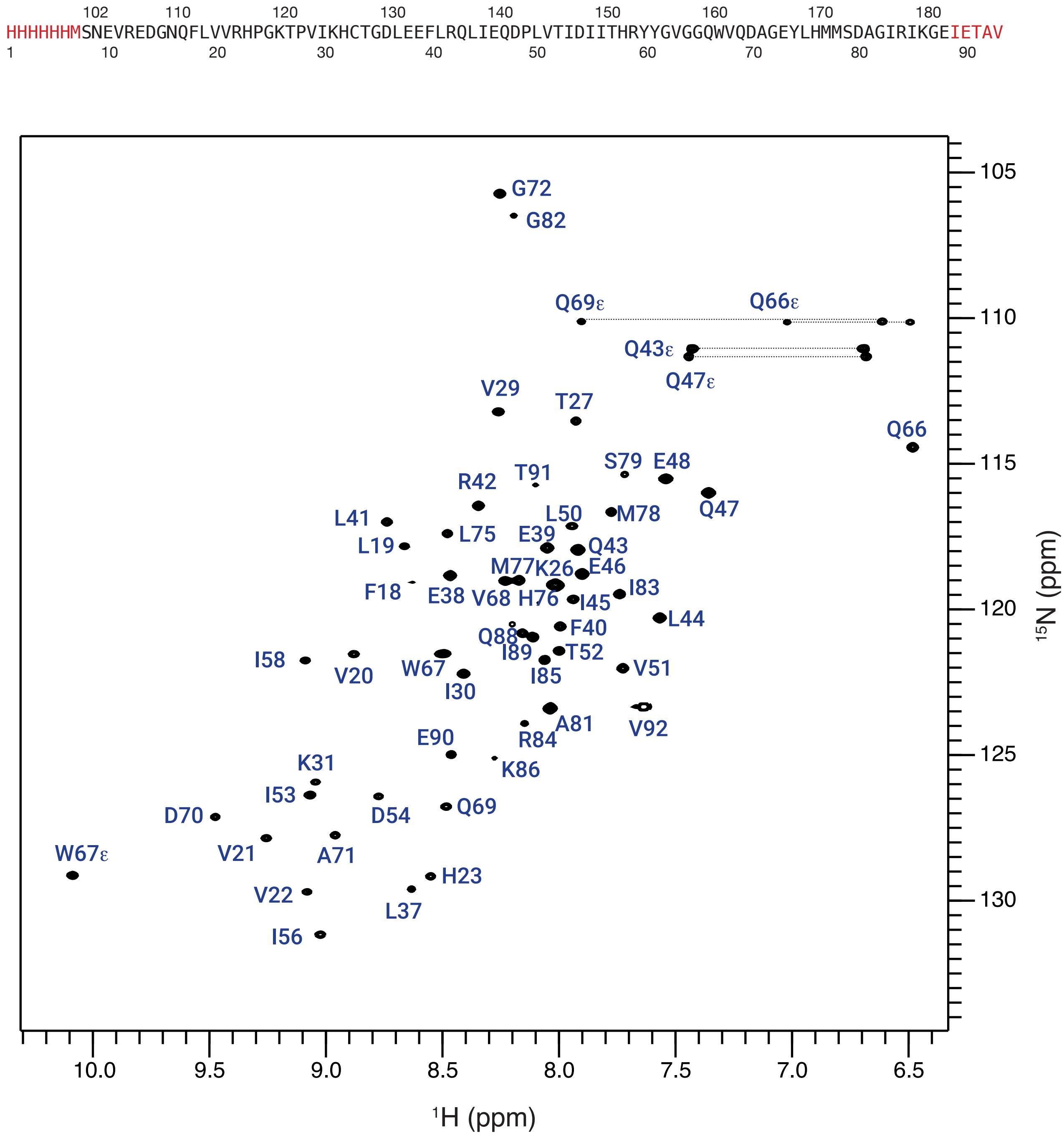


**Figure S1** — A representative $^{1N}$N-edited HSQC spectrum of the Ea22 C-terminal domain. Numbering follows the amino acids in the expressed protein. The panel above translates this numbering scheme to the native Ea22 sequence. Non-native amino acids introduced by cloning are colored red.

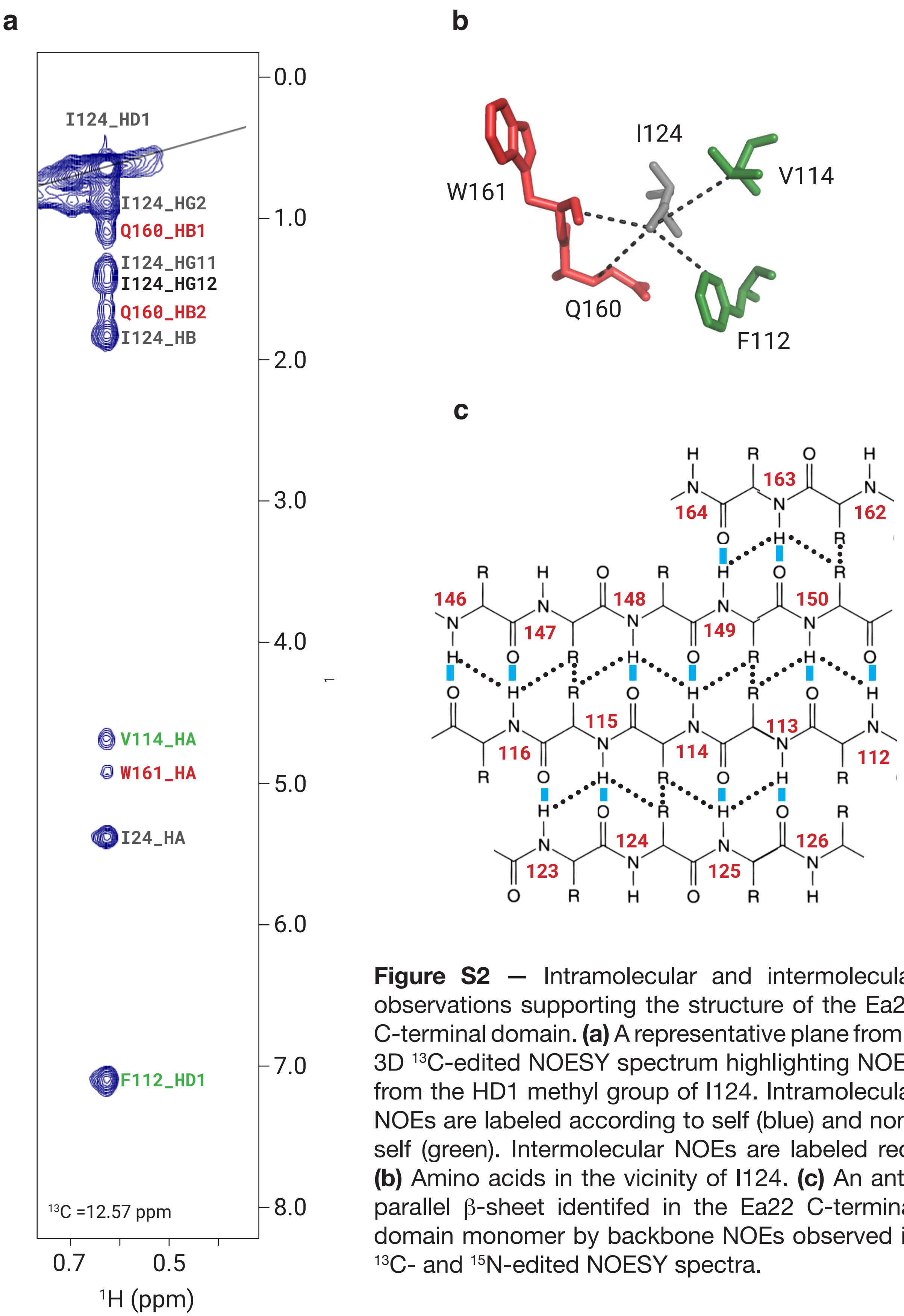


**Figure S2** — Intramolecular and intermolecular observations supporting the structure of the Ea22 C-terminal domain. **(a)** A representative plane from a 3D $^{13}$C-edited NOESY spectrum highlighting NOEs from the HD1 methyl group of I124. Intramolecular NOEs are labeled according to self (blue) and non-self (green). Intermolecular NOEs are labeled red. **(b)** Amino acids in the vicinity of I124. **(c)** An anti-parallel β-sheet identifed in the Ea22 C-terminal domain monomer by backbone NOEs observed in $^{13}$C- and $^{15}$N-edited NOESY spectra.

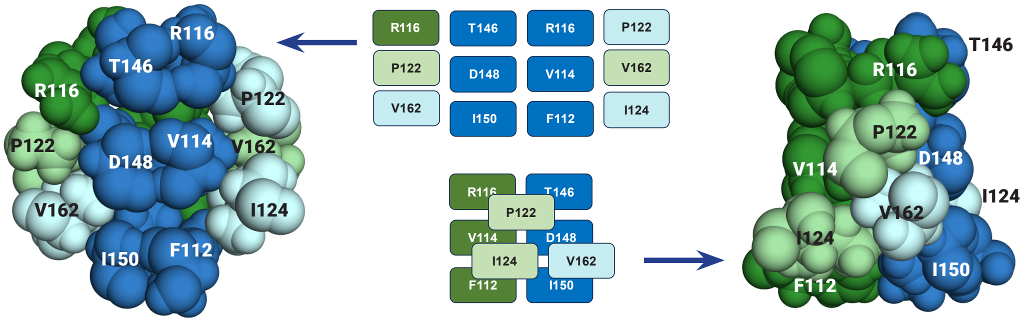


**Figure S3** — Intermolecular interface betwen the ea22 protomers (blue and green shading). Two views of the critical residues are shown, rotated 90°.

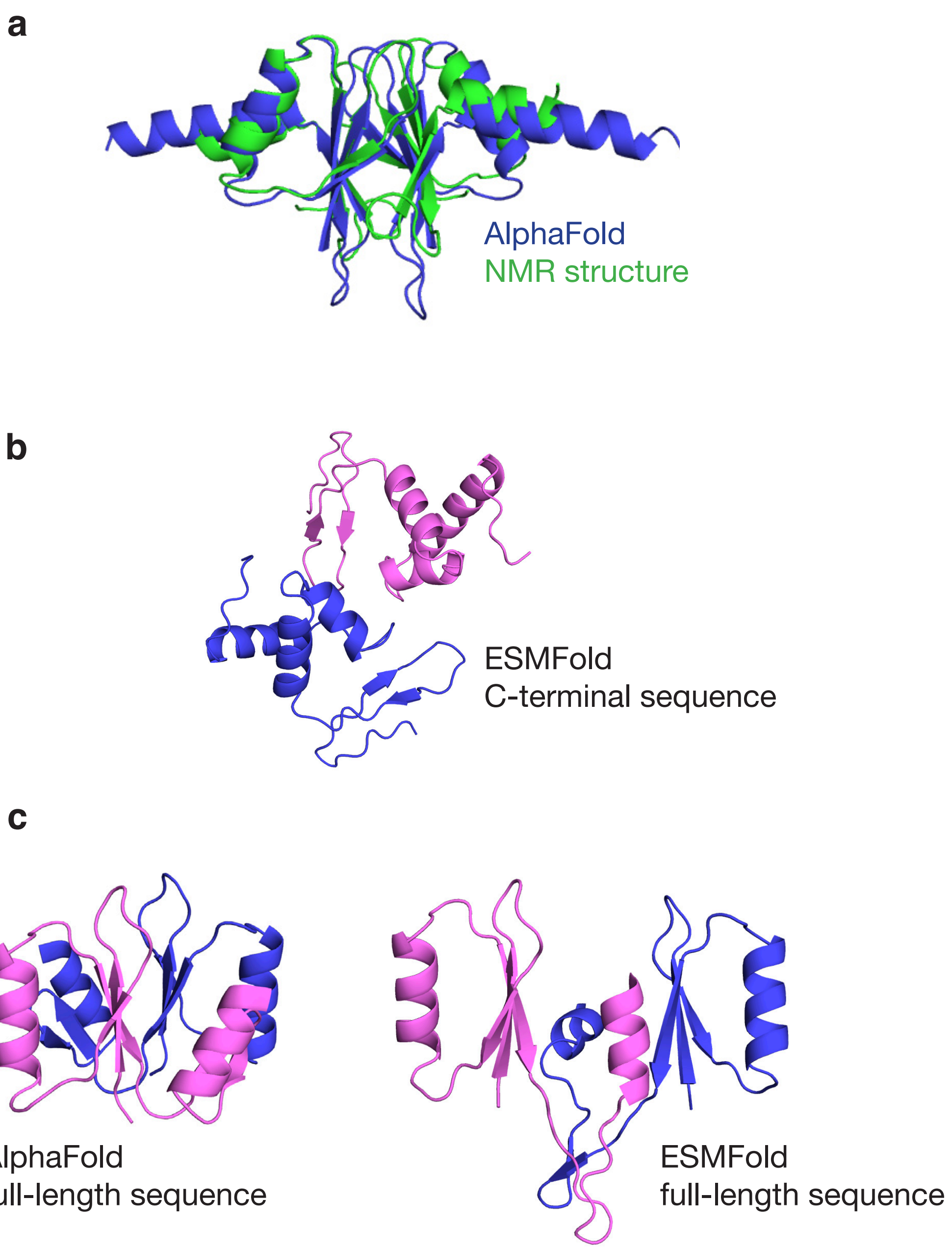


**Figure S4** — Structural predictions of the λ Ea22 CTD dimer. **(a)** The AlphaFold prediction of the C-terminal sequence superimposed with the NMR solution structure. **(b)** A prediction by ESMFold of the C-terminal sequence. **(c)** A prediction by AlphaFold and ESMFold of the full-length Ea22 sequence.

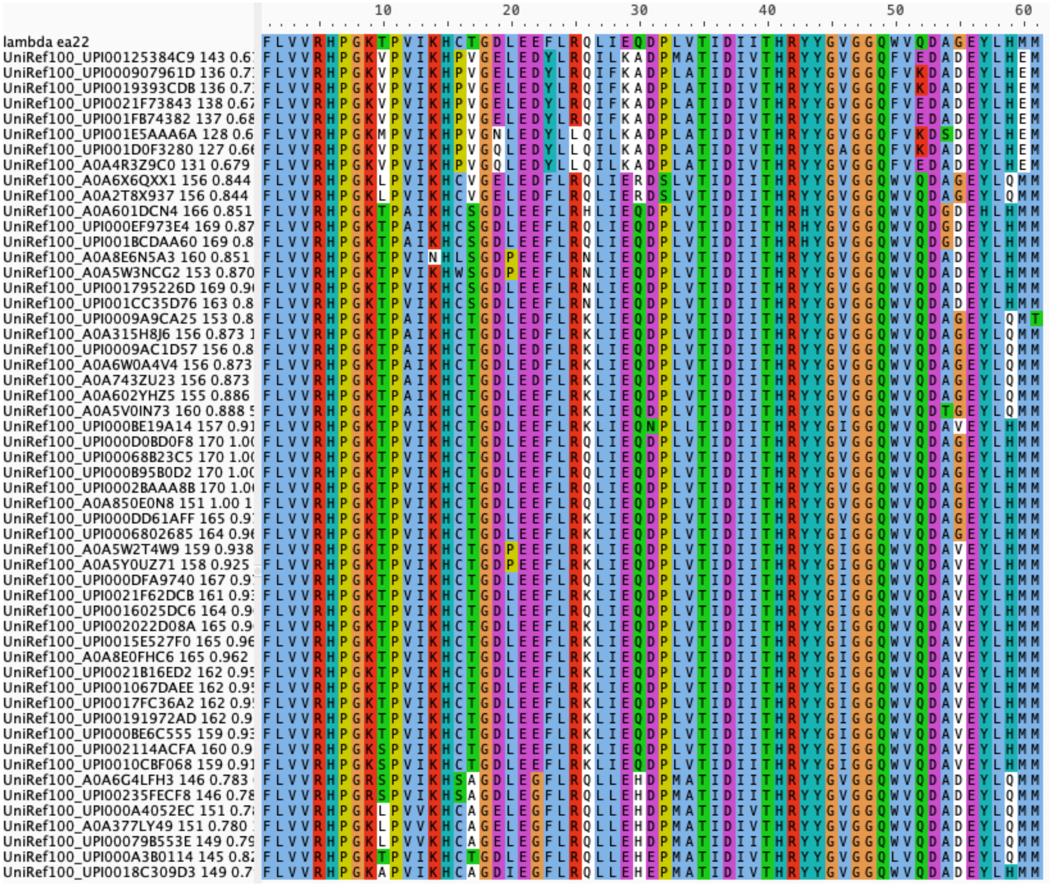

**Figure S5** — Sequence alignment of the λ Ea22 carboxy terminal domain (CTD) and a set of close homologs identified in the UniRef100 database.

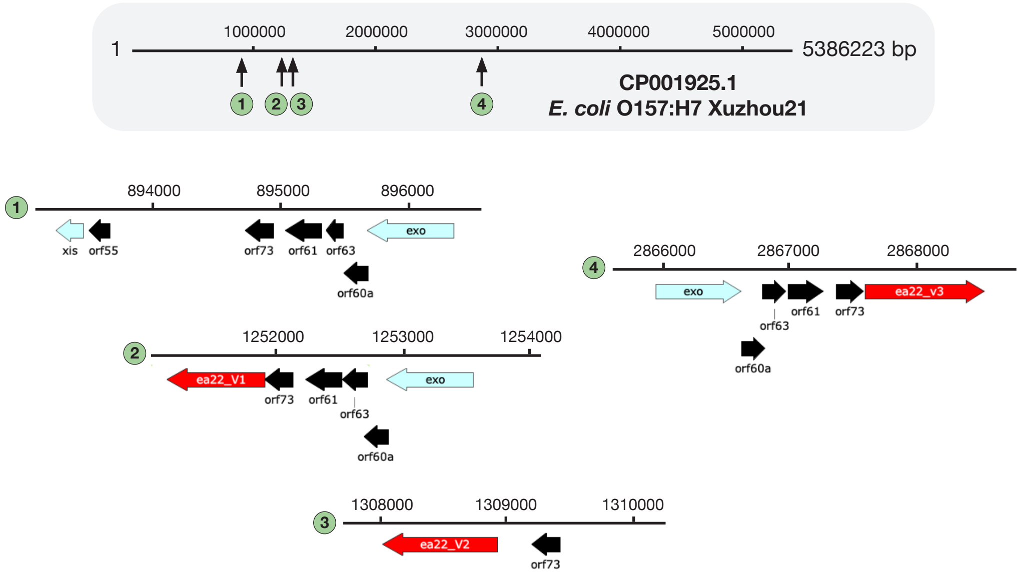


**Figure S6** — Four *exo-xis* gene containing regions are present in the genome of *E. coli* O157:57 str. Xuzhou21.

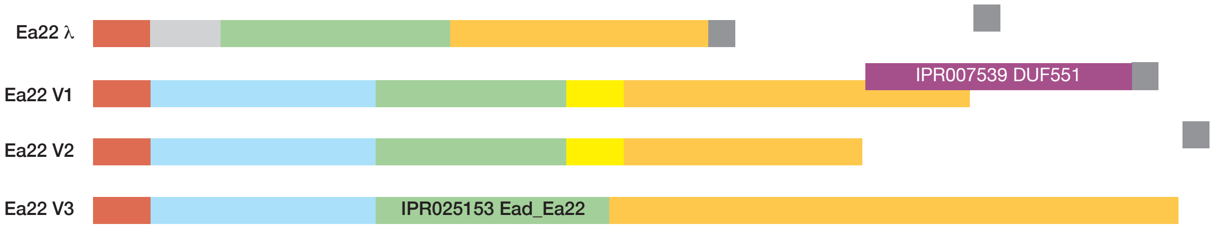


```
lambda    MSEINSQALREAAEQA
V1        MSEINYQALREKAEKA
V2        MSEINYQALREKAEKA
V3        MSEINYQALREKAEKA

lambda    MHDDWGFDADLFHELVTPSI--------------------------------------------
V1        TKGSYIVGHTSVNQHGNLTGVFVCQKWKGEPGGVIAECHVNCLIESDAQAYANAEFIAEANPAT
V2        TKGSYIVGHTSVNQHGNLTGVFVCQKWKGEPGGVIAECHVNCLIESDDQAYANAEFIAEANPAT
V3        TKGSYIVGHTSVNQHGNLTGVFVCQKWKGEPGGVIAECHVNCLVETDAQAYANAEFIAAFNPNV

lambda    VLELLDERERNQQYIKRRDQENEDIALTVGKLRVELETAKSKLNEQREYYEGVISDGSKRIAKLE
V1        VLELLDERERNQQYIKRRDQENEGIALTVGKLRVELEAAENNL-----------IDSECHVAELE
V2        VLALLDEQERNQQYIKRRDQENEEIALTVGKLRVELGAAENNL-----------IDSECHVAELE
V3        ALALLDERERNQQYIKRRDQENEDIALTVGKLRVELEAAKSKLNEQREYYEGVISDGSKHIAELE

lambda    SNEVREDGNQFLVVRHPGKTPVIKHCTGDLEEFLRQLIEQDPLVTIDIITHRYYGVGGQWVQDAGEYLHMMSD
V1        EALRDKLALLEASEKRNAKLQSENAYIRNRYKELDLLIGKNILVMQAAIIEWQATGDAKSGLAWIYNTLFGPG
V2        EALRDKQALLEASEKRIAELEAELVSQTYKLPHTQFEQIANLYEMQFDDGRTCAFHTDAQKAEQWLQACDGNR
V3        KQCAEWERKALSNFEECAAMAERIEEMQTKSAPDSFGIIGENIRTQDNRITSDPMFCVYQKREIVVDADYDYD

lambda    -----------------------------------------------------------------------------------------
V1        ELPDESEKDAQAYFNRKYAPIDEKLMALHKWFWEQSEAKRA------------------------------------------------
V2        VQEYVKLERLQNALSGNSPVTPDGWISCSERMPDTKTAVLVAVEFDRKGDWRMKWATYIPGHPDANDGWIIPGASWKPSHWMPLPEPP-
V3        RIVWVDEDGNEANKLQSRRLELLHENFREPPEKWRRVAVKDIDEFVTCCFTEQGCKDYLAVNGHNLRLPFIYVKSGFRNAEYIGIRNWL

lambda    AGIRIKGE
V1        AGIRIKGE
V2        QEVN-RG-
V3        AGIRIKGE
```

**Figure S7** — Sequence comparison of λ Ea22 and three Ea22 variant prototypes determined from a survey of prophage and phage genomes. The V1 and V3 variants contain regions that are cataloged in the InterPro database.

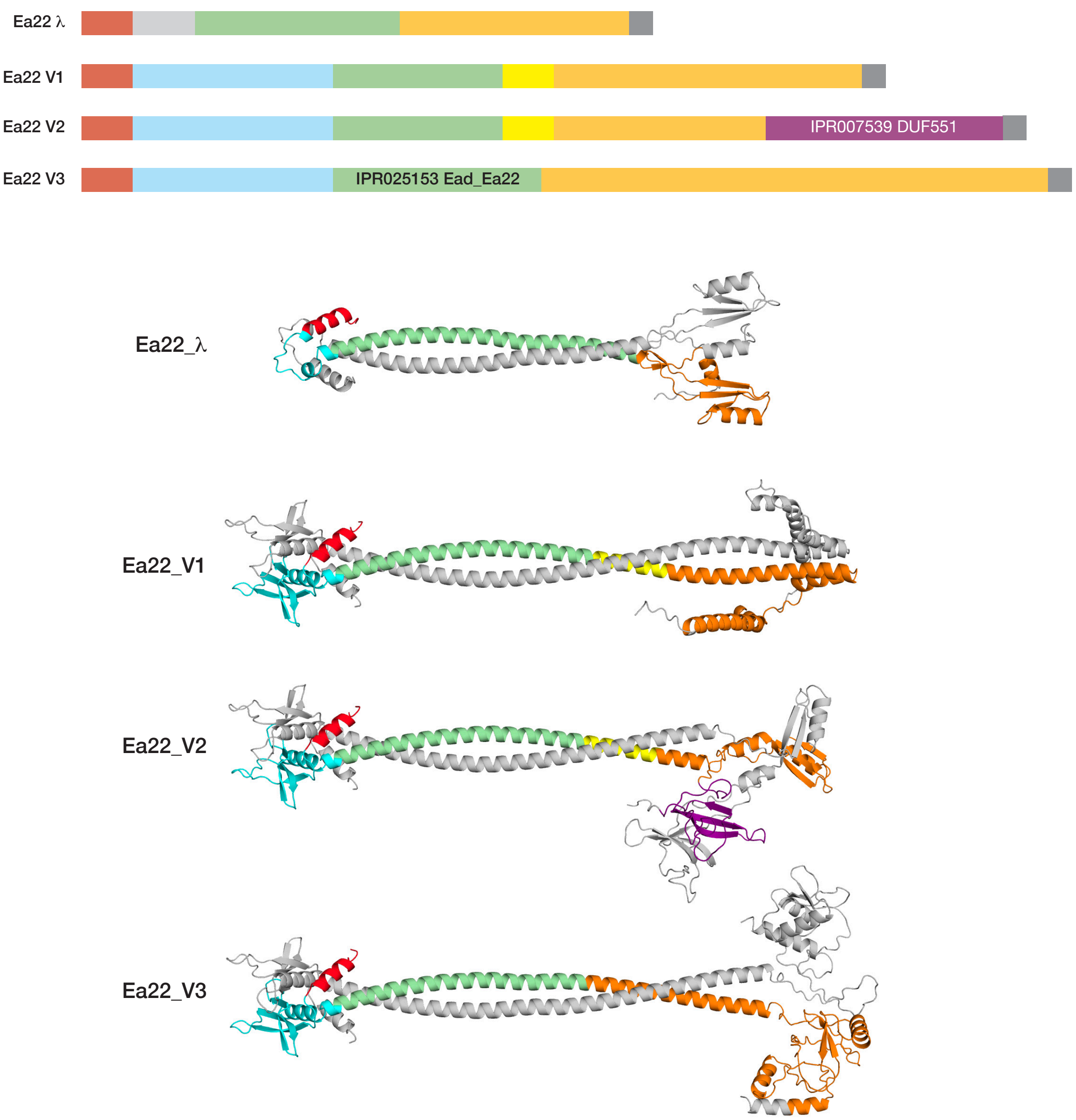


**Figure S8** — ESMFold structure predictions of λ Ea22 and three Ea22 variant prototypes.
